# Biotransformation of lindane (γ-hexachlorocyclohexane) to non-toxic end products by sequential treatment with three mixed anaerobic microbial cultures

**DOI:** 10.1101/2020.10.25.354597

**Authors:** Luz A. Puentes Jácome, Line Lomheim, Sarra Gaspard, Elizabeth A. Edwards

## Abstract

The γ isomer of hexachlorocyclohexane (HCH), also known as lindane, is a carcinogenic persistent organic pollutant. Lindane was used worldwide as an agricultural insecticide. Legacy soil and groundwater contamination with lindane and other HCH isomers is still a big concern. The biotic reductive dechlorination of HCH to non-desirable and toxic lower chlorinated compounds such as monochlorobenzene (MCB) and benzene, among others, has been broadly documented. Here, we demonstrate for the first time that complete biotransformation of lindane to non-toxic end products is attainable using a sequential treatment approach with three mixed anaerobic microbial cultures referred to as culture I, II, and III. Biaugmentation with culture I achieved dechlorination of lindane to MCB and benzene. Culture II was able to dechlorinate MCB to benzene, and finally, culture III carried out methanogenic benzene degradation. Distinct *Dehalobacter* populations, corresponding to different 16S rRNA amplicon sequence variants in culture I and culture II, were responsible for lindane and MCB dechlorination, respectively. This study continues to highlight key roles of *Dehalobacter* spp. as chlorobenzene- and HCH-organohalide-respiring bacteria and demonstrates that sequential treatment with specialized anaerobic cultures may be explored at field sites in order to address legacy soil and groundwater contamination with HCH.

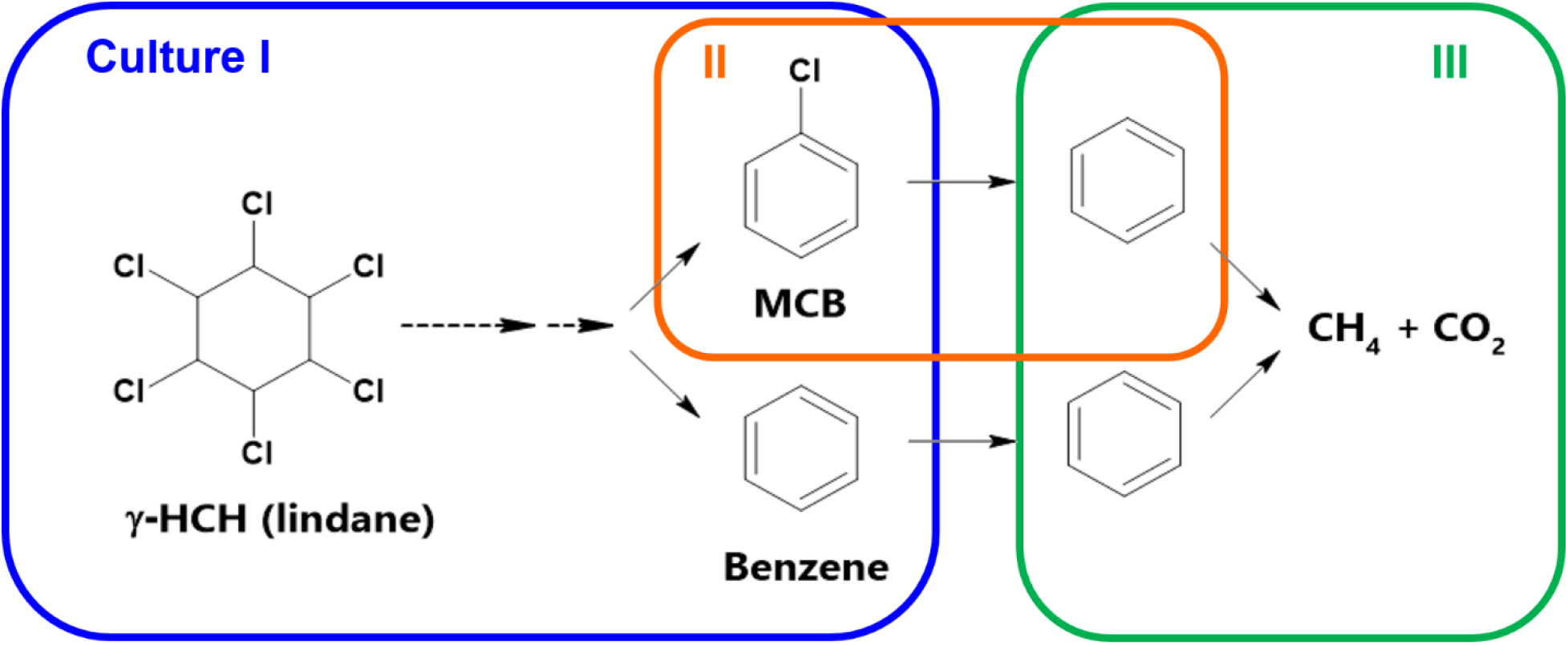

## Introduction

Hexachlorocyclohexane (HCH) isomers are persistent organic pollutants. Lindane, the γ isomer of hexachloroyclohexane (γ-HCH), is the only HCH isomer exhibiting insecticidal properties, and it is a known carcinogen. During the manufacture of “pure” lindane, large amounts of HCH isomers were produced as waste by-products. Lindane (γ-HCH) and other HCH isomers (α-, β-, δ-, and ε-HCH) were components of the agriproduct known as “technical HCH”. Technical HCH and lindane were extensively applied to a variety of crops for pest control. In some instances, HCH manufacture by-products were also improperly disposed of in landfills or waste dumps. Although most countries have banned the use of HCH, groundwater, soil, and sediment contaminated with HCH is of worldwide concern. There is extensive evidence of HCH contamination at a variety of sites in Spain,^1, 2^ Germany,^3^ China,^4^ and the French West Indies,^5^ among others. Legacy HCH contamination, estimated at ~ 4 to 7 million tons,^6–8^ has been difficult to address and manage, likely because of the extent of the contamination and the physicochemical characteristics of each HCH isomer.

Hexachlorocyclohexane isomers are poorly soluble in water. Estimates of water solubility for γ-HCH (lindane) vary; some sources report it to be around 1 to 2 mg/L^9^ and others estimate it to be as high as 7 to 10 mg/L^10, 11^ at temperatures near 20°C. HCH isomers partition into organic compounds (log K_ow_ ~ 3.3) and organic carbon (log K_oc_ ~ 3.0).^12^ Thus, HCHs sorb readily to sediment and soil fractions. HCHs are persistent in the environment, yet the amount of detectable HCH that remains in agricultural soils varies largely depending on the characteristics of the contaminated site; for a review on the persistence of HCH isomers refer to Bhatt et al. (2009).^13^ Long-range global atmospheric transport and deposition of HCHs has also been demonstrated,^14, 15^ resulting in pollution of pristine environments including the Canadian Artic. Natural attenuation of HCH contamination over years or decades has been documented; a summary of these studies is also included in Bhatt et al. (2009).^13^ In light of the extent of HCH contamination and the associated risk posed to human health and the environment, active remediation of contaminated sites may be desirable. Bioremediation could be a suitable approach for HCH transformation as demonstrated at a field study in the Netherlands.^16^

The aerobic biodegradation of HCH is well documented: pathways, microorganisms, and enzymes are known.^17^ The anaerobic biodegradation of HCH has been studied to a lesser extent, yet anaerobic conditions often prevail in HCH-impacted sediments and groundwater. In such environments, HCH isomers can undergo biologically-mediated dechlorination reactions. Dechlorination of HCH isomers has been observed in sediment microcosms,^18^ co-cultures,^19^ anaerobic sludge,^20^ and enrichment cultures.^21^ Known dechlorinating bacteria, such as *Dehalobacter*, *Dehalococcoides,* and *Clostridium* have been linked to HCH dechlorination,^19, 20, 22, 23^ *Dehalobacter* sp. E1 can metabolically dechlorinate β-HCH to MCB and benzene.^19^ Partial HCH dechlorination (in which only a fraction of the added HCH mass was dechlorinated) was observed in pure cultures of *Dehalococcoides mccartyi* strains BTF08 and 195 after being pregrown with tetrachloroethene (PCE);^23, 24^ it has been suggested that *Dehalococcoides mccartyi* strain 195 may be able to respire γ-HCH.^23^ Monochlorobenzene (MCB) and benzene are frequently reported as stable end products of HCH dechlorination. However, since MCB is toxic and benzene is a known carcinogen, these are undesirable biotransformation products. Although MCB and benzene can be biodegraded aerobically, they are often found at contaminated sites along with HCH.^1, 3, 16^ HCH bioremediation could be feasible as long as any potentially toxic byproducts can be biodegraded as well. At some sites, anaerobic biodegradation of these dechlorination by-products may be challenging. However, the sustained anaerobic conversion of MCB and benzene was previously demonstrated in laboratory microcosm experiments^25^ and could be coupled with HCH-dechlorinating bioremediation technologies to achieve complete detoxification.

MCB and benzene, the products of anaerobic HCH dechlorination, have been successfully biotransformed anaerobically.^26–30^ Independent long term investigations in our laboratory have led to the development of three distinct sets of enrichment cultures of interest for HCH bioremediation. The first set of cultures are HCH-dechlorinating enrichment cultures (dechlorinating α-, β-, δ-, and γ-HCH), derived from contaminated sediments from Guadeloupe (French West Indies), which completely transform HCH to MCB and benzene.^21^ Here, the culture that dechlorinates γ-HCH will be referred as culture I. The second culture, or culture II, is a KB-1-derived enrichment culture that dechlorinates MCB to benzene.^26^ The third culture, or culture III, is a methanogenic enrichment culture in which benzene is converted to methane and carbon dioxide^29^ that has been enriched in the Edwards lab for over 20 years. We hypothesized that a combined treatment using the three cultures would be able to completely transform HCH to non-toxic products. Rather than simply mixing the 3 specific cultures together, we tested the hypothesis via sequential addition of each culture. Sequential addition would allow us to minimize any detrimental interactions between the different microbial populations, i.e. the benzene-degrading culture (culture III) uses benzene as electron donor and its activity is impaired by the presence of alternative electron donors which are required for cultures I and II. Using the sequential approach, we were able to demonstrate the anaerobic bioconversion of lindane (γ-HCH) all the way to non-toxic end products. In addition, we demonstrate that distinct native *Dehalobacter* populations, corresponding to different 16S rRNA amplicon sequence variants in culture I and culture II, are responsible for the biotransformation of lindane to MCB and benzene in culture I, and for the dechlorination of MCB to benzene in culture II. Here, we also discuss the implications of our findings which together support the development of anaerobic bioremediation technologies for HCH, MCB, and benzene.

## Materials and Methods

### Chemicals

Chemicals were purchased through Millipore Sigma (Canada) at the highest purity available. Ethanol and methanol for the preparation of feedstocks were HPLC grade. Gases for anaerobic culturing were purchased through Praxair (Canada).

### Enrichment cultures used in this study

Culture I is an anaerobic mixed microbial culture, enriched from soil and river sediment microcosms originating from lindane-contaminated sites in Guadeloupe (French West Indies).^21^ The microcosms were first set up in 2010, and details of the history of culture enrichment are shown in the Supplementary Information (Figure S1). Culture I dechlorinates lindane (γ-HCH, electron acceptor) stoichiometrically to a mixture of MCB and benzene (see Figure 1, panel A, and Figure S2). Ethanol is currently supplied as the electron donor at a ratio of 5 to 1 electron equivalents of donor to acceptor.

**Figure 1.**
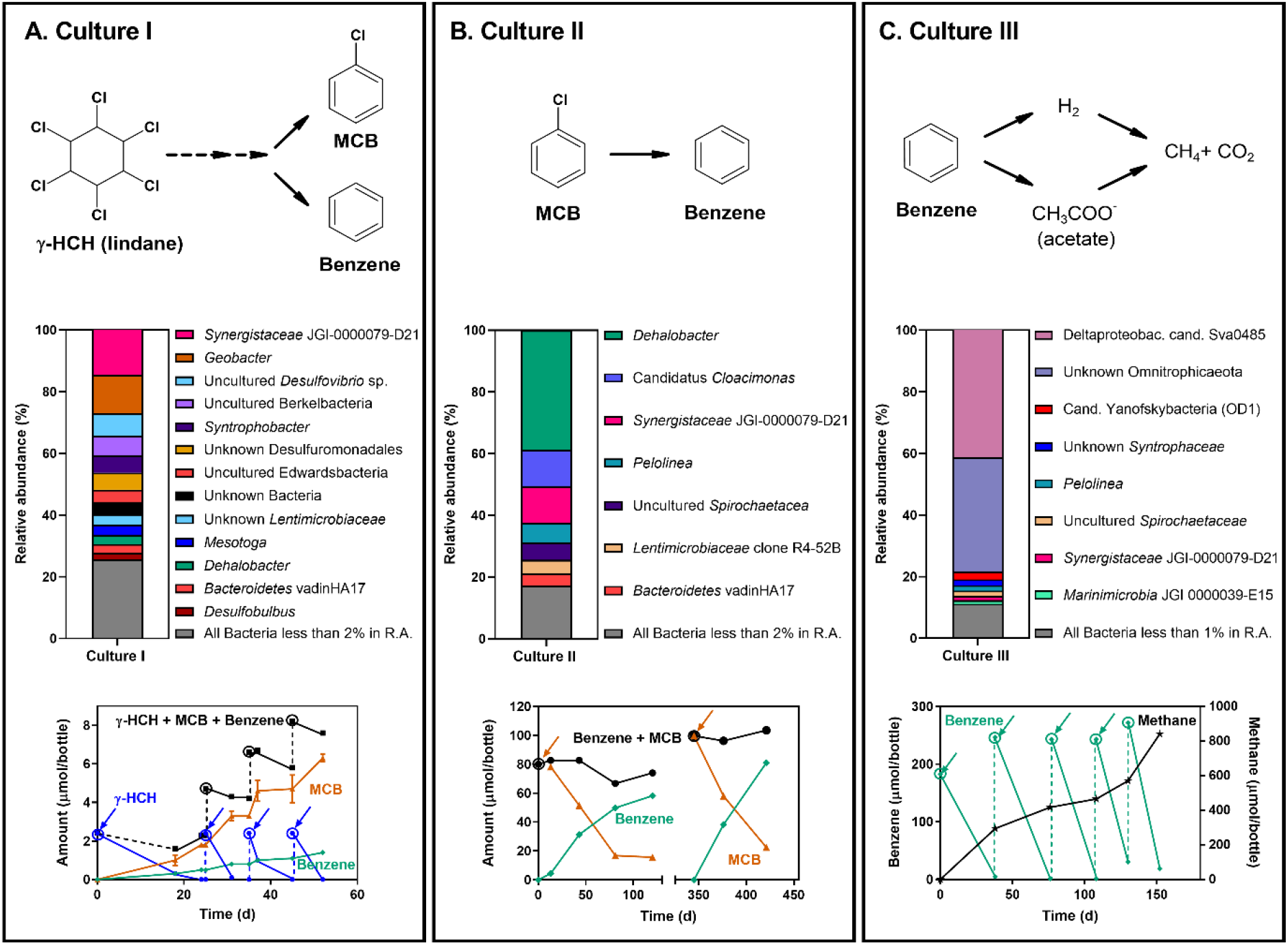
Characteristics of the three microbial enrichment cultures used in this study. Each panel (A,B,C) illustrates the observed biotransformation step (top), the most abundant bacterial groups as determined by 16S rRNA gene amplicon sequencing (middle bar charts), and representative biotransformation data (bottom graph), for each culture. Some bacterial groups could not be specified to genus or species level. These groups were labelled to their most specific level and marked with “unknown”, i.e. “unknown Desulfuromonadales”. Bacterial groups in the bar chart legend are shown as they appear in the bars from top to bottom. In the bottom graphs, arrows indicate feeding events, and circles indicate theoretical and not measured amounts. N.A. stands for not analyzed; * indicates a statistically significant difference when compared to the previous time point, p < 0.05.

Culture II is a KB-1-derived anaerobic enrichment culture that dechlorinates MCB (electron acceptor) to benzene (see Figure 1, panel B). The culture was derived from an enrichment culture that dechlorinates 1,2,4-trichlorobenzene to a mixture of dichlorobenzene, MCB, and benzene.^26^ Culture II is supplied with methanol (electron donor) at a ratio of 5 to 1 electron equivalents of donor to acceptor, but additional re-amendments of donor are generally needed for complete dechlorination of MCB to benzene.

Culture III is an anaerobic methanogenic enrichment culture that transforms benzene (electron donor) through a series of syntrophic microbial interactions to methane and carbon dioxide (see Figure 1, panel C).^29^ This culture is currently being scaled-up at the SiREM Lab (https://www.siremlab.com/) as DGG-B^®^ and pilot tests are underway to evaluate its effectiveness at field scale.

### Sequential treatment with specialized anaerobic microbial enrichment cultures

The biotransformation experiment was performed in three phases, each phase initiated by the addition of one of the three distinct cultures described above (see illustration of experiment setup in Figure S3). A bicarbonate-buffered mineral medium^26^, pre-reduced with an amorphous iron (II) sulfide (FeS) slurry (~0.8 g/L), was used to set up and top up the experimental bottles. All the steps involving feeding, inoculation, and sampling were performed in an anaerobic chamber (Coy Lab Products, USA). All the experiments were set up using 115 mL clear glass bottles with screw cap Mininert^®^ valves to a final liquid volume of 90 mL. During phase I (−150 days), nine active bottles (labelled as experimental bottles 1 to 9 in the supplementary Excel file) containing lindane (~ 4.3 μmol or 1.25 mg per bottle) were inoculated (on Day 0) with a 10% transfer of the lindane-dechlorinating culture (culture I), resulting in a final concentration of 1 x 10^7^ bacterial cells/mL in each experimental bottle. Three additional bottles were inoculated with 10% autoclaved inoculum (negative controls); the inoculum was autoclaved in a liquid cycle held at 121°C and 15 psi for 45 min. Glass bottles were pre-loaded with lindane prior to bringing them into the anaerobic chamber as follows: lindane was added to autoclaved empty bottles as a concentrated lindane/acetone solution which was completely evaporated under a stream of nitrogen. The bottles were then transferred to the anaerobic chamber for a minimum of 2 days prior to inoculation. During inoculation, sterile mineral medium and culture I were aliquoted directly into each bottle using sterile pipettes. On Day 93, after all lindane had been converted to MCB and benzene, cultures were transferred into new bottles with additional lindane (~ 8.6 μmol or 2.25 mg per bottle, prepared as described above). Electron donor, ethanol, was added every four weeks to experimental and control bottles (on Day 0, 27, 59, 93, and 123) at a concentration of 0.3 mM. The total amount of donor added during phase I is equivalent to 20 times the total electron equivalents of acceptor added (assuming 6 electron equivalents per mole of lindane).

During phase II, from Day 148 to 337 (−190 days), six of the nine active bottles from phase I (labelled as experimental bottles 1 to 6 in the supplementary Excel file) were bioaugmented on two separate dates (Day 148 and Day 275) with the MCB-dechlorinating culture (culture II). For each bioaugmentation event of phase II, the experimental bottles were inoculated with 1 x 10^6^ bacterial cells of culture II per mL. From phase II onwards, cultures were added to each bottle as concentrated inoculum using sterile plastic syringes via the septum on the screw cap to minimize the potential loss of MCB and benzene and to replenish the experimental working volume (90 mL). For the first and second phase-II bioaugmentation events, 5 mL of 10X-concentrated culture II and 10 mL of 4X-concentrated culture II were added to each bottle, respectively (refer to the SI for concentration protocol). Culture II-bioaugmented bottles received methanol as electron donor. Methanol was fed at a concentration of 0.1 mM (Day 148 and Day 198) and 0.2 mM (Day 218, 258, 275). The total amount of methanol added to these bottles throughout phase II was equivalent to 50 times the electron equivalents of acceptor (total MCB produced during the experiment; 2 electron equivalents per mole of MCB). The three remaining active bottles from phase I were not inoculated with culture II, but were monitored as controls. Ethanol was added to these control bottles on Day 198, 218, and 296.

During phase III (Day 148 to 337), three of the six active bottles that were bioaugmented in phase II (labelled as experimental bottles 4, 5, and 6 in the supplementary Excel file) were inoculated with 4 x 10^8^ bacterial cells of the benzene-degrading culture (culture III) per mL, while the remaining three bottles were monitored as controls. Additionally, two positive controls, inoculated with culture III in the same fashion as bottles 4, 5, and 6, were set up in parallel to monitor the activity of the benzene-degrading culture; neat benzene was added directly to fresh mineral medium. Similarly to phase II, culture III was added to each bioaugmented bottle as concentrated inoculum (6 mL at 6X) using sterile plastic syringes via the septum on the screw cap (refer to the SI for concentration protocol).

### Analytical methods

MCB, benzene, and methane were quantified using GC-FID (gas chromatography with flame ionization detection). For the analysis, aqueous samples (1 mL) were added to 5 mL of acidified (using 6N HCl) deionized water (pH < 2). Calibration standards for MCB and benzene were prepared gravimetrically by adding a known mass of a combined MCB/benzene stock prepared in methanol to bottles containing known volumes of water and headspace. Methane standards were prepared by adding known volumes of pure methane gas, using gas-tight syringes, to bottles containing known volumes of water and headspace. External calibrations were prepared to convert area counts into concentrations (refer to the SI for details on GC method). Analysis of γ-hexachlorocyclohexane (lindane) was not performed for the sequential biotransformation experiment since that would have required removal of more than 1 mL of culture volume per bottle per sampling point during phase I. However, lindane was measured in a separate experiment shown in Figure S2, in which it was verified that culture I biotransforms lindane to MCB and benzene stoichiometrically (moles of lindane degraded = moles of MCB + benzene produced). This stoichiometric relationship between lindane fed and MCB and benzene produced has consistently been observed in this culture throughout its enrichment process.

### DNA sampling and quantitative PCR (qPCR)

DNA was extracted from 2 mL culture samples. Cells were harvested by centrifugation at 10,000x g for 15 min at 4°C. Cell pellets were resuspended in 100 μl of supernatant and DNA was extracted using the DNeasy PowerSoil kit following the manufacturer’s recommendations (during steps 9 and 12 of the protocol, all supernatant was transferred). Real-time polymerase chain reaction (qPCR) assays were performed to track the gene copies of total Bacteria, *Dehalobacter* and Desulforomonadales using specific 16S rRNA gene primers (see Table S1 and Table S2 in the SI for qPCR primers and calibration information, respectively). For this study, a new primer set that targets representative sequences classified as either Desulforomonadales and/or *Geobacter* (a genus of the order Desulforomonadales) was designed; the target sequences were obtained from previous 16S rRNA gene amplicon sequencing studies of culture I^21^ (refer to the SI for the qPCR prep and quantification protocol).

### 16S rRNA gene amplicon sequencing

DNA was extracted as described above and sent for Illumina amplicon sequencing at the Genome Quebec Innovation Centre (McGill University). DNA was sequenced using the Illumina MiSeq PE300 platform with modified versions of primers 926f and 1392r (926f-modified: AAACTYAAAKGAATWGRCGG; 1392r-modified: ACGGGCGGTGWGTRC) which target the V6-V8 variable region of the 16S rRNA gene from bacteria and archaea, as well as the 18S rRNA gene in eukarya. After sequencing, raw data was processed using the QIIME 2^31^ Core 2020.2 distribution. Primers were removed using the cutadapt^32^ plugin. Sequences were denoised and merged into amplicon sequence variants (ASVs) using DADA2^33^ (forward sequence truncated at 260 bp, reverse sequence truncated at 240 bp). Taxonomy was assigned to ASVs using the feature-classifier plugin^34^ with the classify-sklearn method using a pre-trained Naïve Bayes Classifier with the SILVA 132 99% OTUs full-length sequences database (available at https://docs.qiime2.org/2020.2/data-resources/). Data was taken to MS Excel and RStudio for further processing. An Excel supplementary file is included with the resulting taxonomy and summary of ASVs of interest. Raw 16S rRNA amplicon sequencing data has been deposited to the NCBI Sequence Read Archive under bioproject PRJNA661282.

### Estimation of absolute abundance of amplicon sequence variants (ASVs)

Absolute abundance of ASVs of interest was calculated as the product of the qPCR-determined Bacteria 16S rRNA gene copy numbers (determined using the general Bacteria primer shown in Table S1) and the relative abundance of each ASV (included in the Supplementary Excel file) determined by 16S rRNA gene amplicon sequencing via the QIIME 2 pipeline (described above). This approach was used to obtain the absolute abundance of bacterial groups of interest during phase III and for the different *Dehalobacter* ASVs.

## Results and Discussion

### Inoculum samples from enrichment cultures used during the biotransformation experiment

Samples from cultures I, II, and III were used as inoculum for each respective phase of the biotransformation experiment. Figure 1 shows the observed biotransformation step (top diagram), the bacterial community composition determined by 16S rRNA amplicon sequencing at the time of inoculation (middle), and culture maintenance data (bottom) for each culture; cultures I, II, and III, are shown in panels A, B, and C, respectively. For culture I (Figure 1A), lindane is dechlorinated to MCB and benzene. The proposed series of reactions that leads to the formation of MCB and benzene in culture I are discussed in Qiao et al.^21^ Dechlorination rates for this culture range from ~ 6.0 to 13 μmol/L/day (1.8 – 3.8 mg/L/day). Detected abundant bacterial genera (> 2% relative abundance) that are known organohalide-respirers include *Geobacter,* and *Dehalobacter.* Other abundant bacterial genera include representatives of the fermentative and sulfate-reducing genus *Desulfovibrio,* members of the recently revealed candidate phyla Berkelbacteria, as well as members of the *Syntrophobacter,* and the Desulforomonadales family. Culture II, which dechlorinates MCB to benzene, contains abundant bacteria classified as *Dehalobacter, Cloacimonas*, *Synergistaceae, Pelolinea,* and *Spirochaetacea* (Figure 1B). MCB dechlorination to benzene is observed a rate of ~ 2.0 ± 0.3 μmol/L/day (0.16 ± 0.02 mg/L/day). Culture III, degrades benzene to methane and carbon dioxide (Figure 1C). Culture III is highly enriched in bacteria classified as Deltaproteobacterial candidate Sva0485 which represents the key microorganism, historically termed “ORM2” for Oil Refinery Methanogenic Clone 2, responsible for the initial attack on the benzene ring.^30, 35^ Other abundant bacteria in this culture belong to the phyla Omnitrophicaeota and Yanofskybacteria. The ASVs classified as Yanofskybacteria correspond to another previously described abundant bacterium in the benzene-degrading cultures, referred to as OD1. ^29^ The benzene degradation rate for culture III used in this study was ~ 6.4 μmol/L/day (~ 0.5 mg/L/d).

### Phase I: dechlorination of lindane (γ-HCH) to MCB and benzene

Figure 2A shows the dechlorination of lindane from Day 0 to Day 173, in the 9 experimental bottles inoculated with culture I. Within 14 days, about 96% of the theoretical mass of lindane added to the bottles was dechlorinated at an average rate of 3.3 ± 0.4 μmol/L/d (1.0 ± 0.1 mg/L/d). By Day 23, all lindane was transformed to MCB and benzene. Even though electron donor (ethanol) was re-amended (Day 27 and Day 59), further dechlorination of MCB to benzene did not occur. This is consistent with the observations gathered for this culture during years of sequential transfers and maintenance (Figure S1, Supplementary Information). For the second feeding event (Day 93), the added mass of lindane was 1.8 times greater than the first feeding. As expected, corresponding stoichiometric amounts of MCB and benzene were produced. After 13 days (Day 106), 53% of the re-amended theoretical mass of lindane was dechlorinated at an average dechlorination rate of 3.9 ± 0.3 μmol/L/d (1.1 ± 0.1 mg/L/d). Electron donor limitation became evident after sampling on Day 121, so ethanol was re-amended (Day 123). All lindane was dechlorinated afterwards. At the end of phase I, the average molar ratio of benzene to MCB was 0.32 ± 0.06. In the control bottles with autoclaved inoculum, only trace amounts of benzene and MCB were detected (Figure 2B), likely due to abiotic reductive dechlorination.

**Figure 2.**
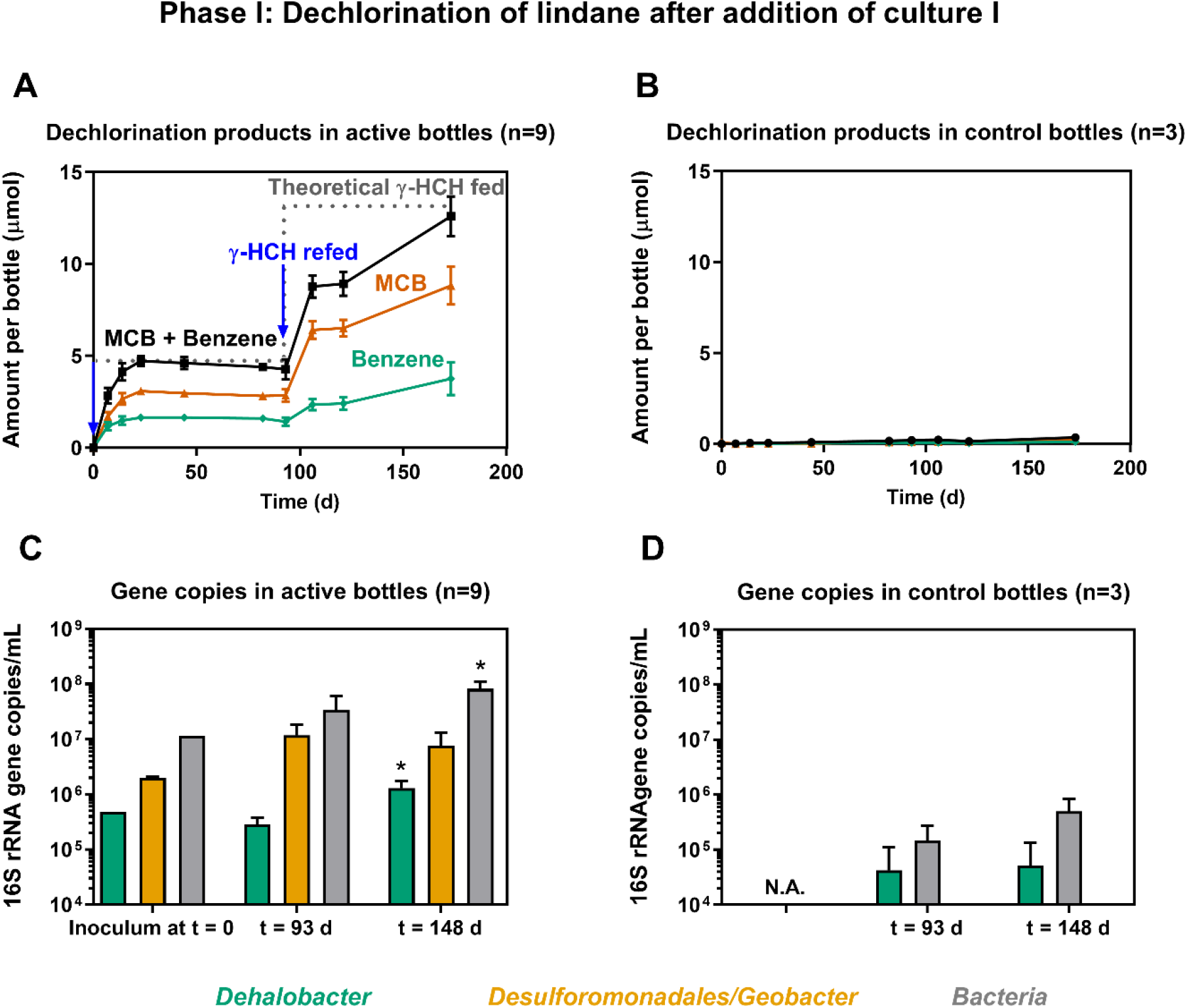
Dechlorination of lindane to monochlorobenzene (MCB) and benzene during Phase I of the lindane biotransformation experiment. The figure shows the concentration of MCB and benzene over time in active (A) and control bottles (B), and the 16S rRNA gene copies per mL of culture of *Dehalobacter*, Desulforomanadales*/Geobacter*, and Bacteria in active (C) and control bottles (D). Culture I was added on Day 0. Error bars represent the standard deviation of replicate experimental bottles (n). Blue arrows indicate lindane feedings; * indicates a statistically significant difference when compared to the previous time point, p < 0.05.

The qPCR-determined 16S rRNA gene copies per mL of culture of *Dehalobacter,* Desulfuromomadales/*Geobacter*, and total Bacteria are shown in Figure 2C and D. There was a significant increase in *Dehalobacter* and total Bacteria copy numbers from Day 93 (prior to the second lindane feeding) to Day 148. *Dehalobacter* copy numbers increased by 4.3-fold. Desulfuromomadales/*Geobacter* copy numbers did not increase. This indicates that the growth of *Dehalobacter* is linked to the dechlorination of lindane to MCB and benzene, and that bacteria belonging to the Desulfuromomadales/*Geobacter* are likely not directly implicated. These findings are consistent with observations reported earlier for culture I.^21^ Since some *Geobacter* spp. are known to dechlorinate organochlorines^36–38^, we had hypothesized that they could be involved in lindane dechlorination. *Geobacter* in culture I are potentially using ethanol (supplied during culture maintenance) or other bacterial metabolites, e.g. acetate or H2 as electron donor. The use of ethanol as electron donor is a known feature of some *Geobacter* spp. such as *G. chapellei* strain 172T, *G. metallireducens* GS-15T, *G. hydrogenophilus* H-2T, and *G. grbiciae* TACP-2T^37, 39^ In addition, *G. hydrogenophilus* H-2T, *G. grbiciae* TACP-2 T, and *G. sulfurreducens* can use use H2 as electron donor^39^ and require acetate as a carbon source for growth with H2. *Geobacter* spp. can use a wide range of electron acceptors, e.g. elemental sulfur, nitrate, malate, Mn (IV), Fe (III), and some organochlorines, and can also transfer electrons directly to other microbes^40^. Bacteria belonging to the Desulfuromomadales could also be fermenting organic acids.

Interestingly, *Dehalobacter* but not Desulfuromomadales/*Geobacter* copy numbers were quantifiable via qPCR in the bottles with autoclaved inoculum. *Dehalobacter* gene copies were also detected in heat-treated experimental control bottles from a previous study with *Dehalobacter*-enriched cultures.^26^ *Dehalobacter* spp. are members of the Firmicutes which are known to produce heat-resistant endospores.^41^ Endospores may survive the autoclaving process and thus result in cell viability and DNA preservation. *Dehalobacter* genomes do contain sporulation genes^42^, yet the *Dehalobacter restrictus* type strain, DSM 9455, formerly known as PER-K23, was reported as not spore-forming.^43^ Additional studies on *Dehalobacter* spp. are required to determine if spore formation is a feature of select strains.

### Phase II: dechlorination of MCB to benzene

On Day 148, six of the active bottles (experimental bottles 1 to 6) from phase I were bioaugmented with culture II. Experimental bottles 7, 8, 9 were not bioaugmented with culture II, and remained as controls. Figure 3 shows the steady dechlorination of MCB to benzene in five out of the six bioaugmented bottles; one of the bioaugmented bottles did not initially exhibit significant MCB dechlorination, and is not included in the graph. A slight reduction in dechlorination rates was observed for the time period between Day 237 and 254 with some bottles exhibiting significant deviations from the mean dechlorination rate. As a result, a second bioaugmentation event, in which culture II was again added to experimental bottles 1 to 6, was performed on Day 275. Interestingly, after this bioaugmentation event, MCB dechlorination was kick-started in the single bottle in which no significant MCB dechlorination had been observed resulting in detectable MCB to benzene transformation in all the bioaugmented bottles. The mean MCB to benzene dechlorination rate for phase III was 0.3 ± 0.2 μmol/L/day (0.03 ± 0.02 mg/L/day, n = 5). This rate is about 5 times lower than the average rate measured during routine cultivation of culture II (0.03 vs 0.16 mg/L/day). One of the factors that may have contributed to the rate reduction is the increased competition for electron donor that the MCB-dechlorinating population likely faced in these experimental bottles. As shown in Figure 1, culture I (lindane enrichment) is a mixed enrichment culture with organisms known to be able to use methanol, e.g. for methane production in certain members of the *Methanosarcinales*, or potentially as electron donor. Several amplicon sequence variants in culture I were classified to the genus *Desulfosporosinus*. Some *Desulfosporosinus* spp. such as *D*. *meridie,* a homoacetogenic bacterium, use methanol as electron donor during sulfate reduction.^44^

**Figure 3.**
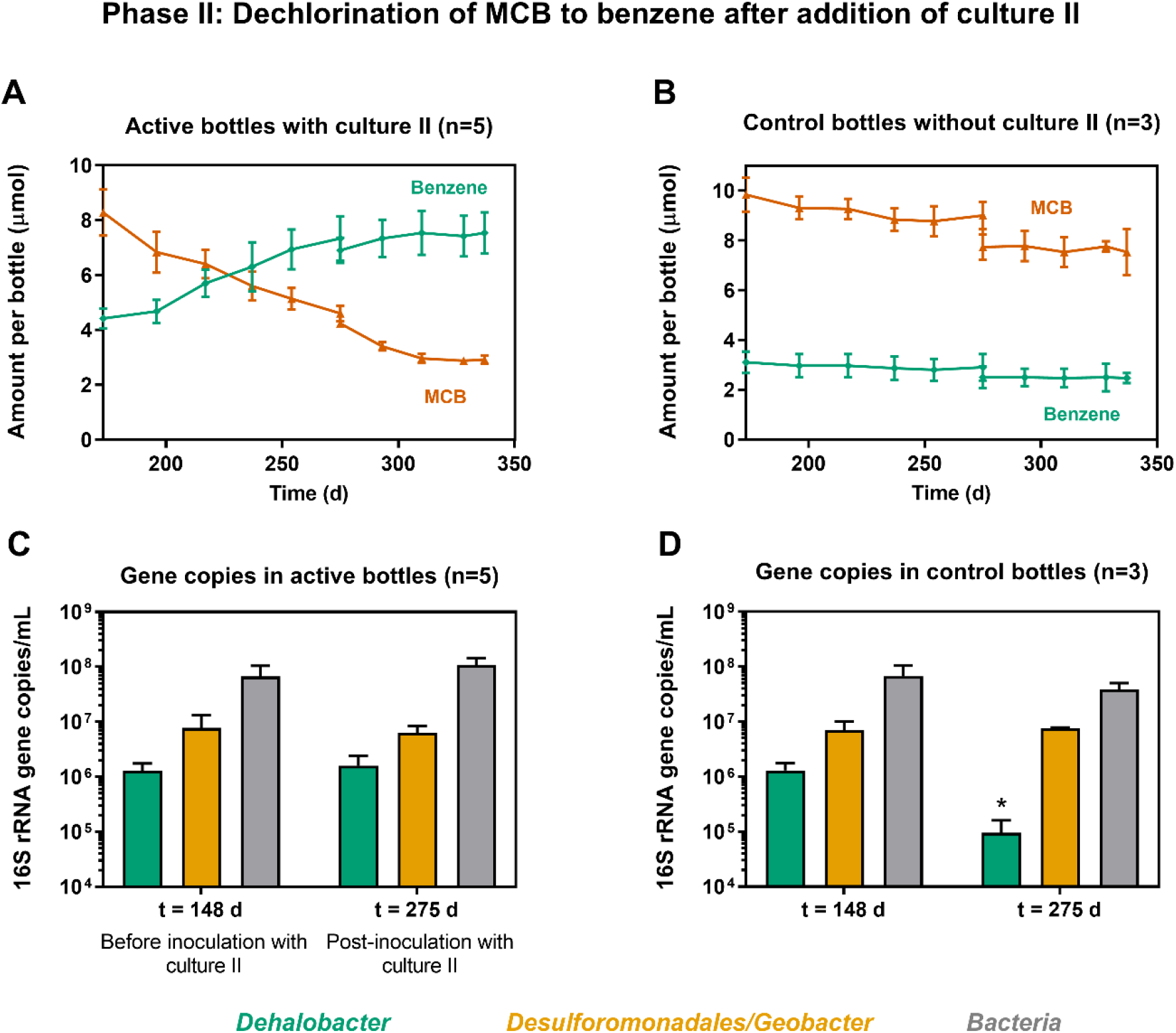
Dechlorination of MCB to benzene during Phase II of the lindane biotransformation experiment. The figure shows the concentrations of MCB and benzene in active (A) and control bottles (B), and 16S rRNA gene copies per mL of culture of *Dehalobacter,* Desulforomanadales*/Geobacter*, and Bacteria in active (C) and control bottles (D). Culture II was added on Day 148 and Day 275. Error bars represent the standard deviation of replicate experimental bottles (n). N.D stands for non-detected. N.A. stands for not analyzed; * indicates a statistically significant difference when compared to the previous time point, p < 0.05.

Benzene and MCB amounts in the control bottles (non-bioaugmented with culture II) are shown in Figure 3B. As expected, benzene did not increase in these bottles. As phase II progressed, there was a slight decline in the amount of MCB and benzene measured in the control and active bottles. This was due to addition of inoculum (10 mL) to the active bottles during the second bioaugmentation event, and to the fact that all the experimental bottles were topped up to 90 mL on Day 275. As shown in Figure 3C, the qPCR-measured Bacteria and *Dehalobacter* gene copies per mL of culture did not increase in the active bottles over the 127-day period following inoculation. However, there was a measurable decline (14-fold) of *Dehalobacter* gene copies per mL of culture in the control (non-bioaugmented) bottles (Figure 3D). This finding provides additional evidence for the role of the *Dehalobacter* population in culture I as the lindane-dechlorinating population. In the active bottles, the *Dehalobacter* gene copies did not decline because the lindane-dechlorinating *Dehalobacter* population was being replaced by the MCB-dechlorinating *Dehalobacter* population native to culture II. This will be further discussed later. In the controls (non-bioaugmented with culture II, Desulfuromomadales/*Geobacter* gene copies did not decline suggesting that these organisms were metabolizing some of the available endogenous electron donor (e.g. acetate) that likely was still present in the bottles; ethanol was added to these controls in two occasions (Day 198 and 218).

On Day 337, before proceeding with the addition of culture III, some MCB still remained in the culture II-bioaugmented bottles. For complete MCB dechlorination to have occurred, additional bioaugmentation events, likely with a higher inoculum density, would have been required. As will be discussed later, the *Dehalobacter* population in culture II is usually in the order of 10^7^ cells/mL of culture, not very dense compared to other dechlorinating enrichment cultures, and its calculated yield is lower than that of other *Dehalobacter* populations respiring different chlorinated substrates.

### Phase III: benzene biodegradation

After 190 days into phase II (Day 337), we proceeded with phase III, i.e. the bioaugmentation with the benzene-degrading culture (culture III). At this point, 67 ± 7% of the mass of MCB had been converted to benzene following the bioaugmentation with culture II. Culture III was added to three of the active bottles from phase II (experimental bottles 4, 5, and 6); the remaining three bottles from phase II served as controls. As shown in Figure 4, benzene was biodegraded at a rate of 1.3 ± 0.1 μmol/L/day (0.1 ± 0.01 mg/L/day) in two out of the three bioaugmented bottles. In the remaining bioaugmented bottle, benzene degradation occurred as well but at a lower rate. In the controls (not bioaugmented with culture III), benzene degradation did not occur (Figure 4B).

**Figure 4.**
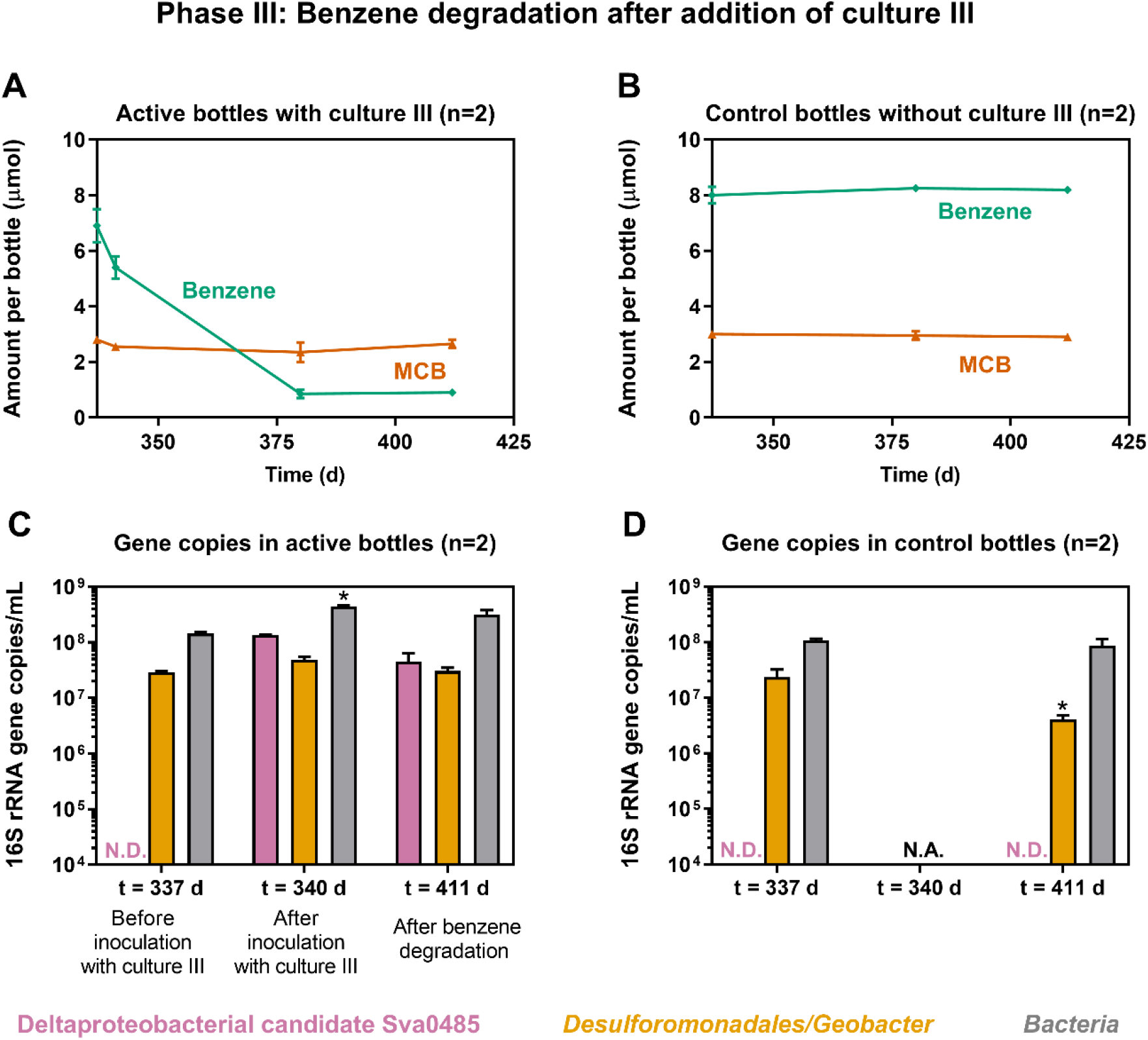
Benzene degradation during Phase III of the lindane biotransformation experiment. The figure shows the concentrations of MCB and benzene in active (A) and control bottles (B), and 16S rRNA gene copies per mL of culture of Deltaproteobacterial candidate Sva0485, Desulforomanadales*/Geobacter*, and Bacteria in active (C) and control bottles (D). Culture III was added on Day 337. Error bars represent the range of measurements in duplicate experimental bottles. N.A. stands for not analyzed. N.D. stands for non-detectable; * indicates a statistically significant difference when compared to the previous time point, p < 0.05.

After the bioaugmentation with culture III, the resulting gene copies per mL of Deltaproteobacterial candidate Sva0485 (previously referred to as ORM2), were in the order of 10^8^ and remained at 10^7^ after benzene degradation (Figure 4C). Recall that ORM2 is the bacterium responsible for the initial attack on the benzene ring. The average amount of benzene produced in the experimental bottles (6.5 mg/L) was likely not enough to maintain the population of ORM2 at such high density (10^8^ ORM2 gene copies per mL); culture III is generally maintained at higher concentrations, ranging from 30 to 150 mg/L, in order to support high cell density. The gene copies per mL for Desulfuromomadales/*Geobacter* remained at 10^7^ for the active bottles and declined to 10^6^ in the control bottles (Figure 4D). In the active bottles, endogenous electron donors may have been available to sustain the population of Desulfuromomadales/*Geobacter*. In the control bottles, electron donor was not added during phase III, and thus the microbial community derived from phase I likely faced electron donor shortage. The decline in Desulfuromomadales/*Geobacter* copy numbers in these control bottles strongly suggests that these organisms were metabolizing some of the available electron donor which was likely exhausted during the transition from phase II to phase III (methanol was last supplied on Day 296).

Neat benzene (not produced from lindane dechlorination) was subsequently added to the three experimental bottles on Day 441 to confirm that the benzene-degrading microbial community remained active. As shown in Figure S4, benzene was indeed biodegraded afterwards. The bottles that were not bioaugmented with culture II or culture III did not exhibit further dechlorination or benzene degradation.

### MCB is dechlorinated to benzene in culture II by a native KB-1-derived *Dehalobacter* population distinct from the native *Dehalobacter* population in culture I

As mentioned earlier, culture II originated from a KB-1-derived enrichment culture (referred to in the lab as 1,2,4TCB/M_2014) that dechlorinates 1,2,4-trichlorobenzene (1,2,4-TCB) to a mixture of dichlorobenzene, MCB, and benzene.^26^ Culture II (500 mL scale-up) had been maintained on MCB as the sole electron acceptor for about 2 years. To clearly demonstrate *Dehalobacter* population growth during MCB dechlorination to benzene, the culture was sampled frequently over a period of 127 days and MCB/Benzene and *Dehalobacter* abundances were analyzed. As shown in Figure S5, during the dechlorination of ~ 100 μmol of MCB, the relative abundance of *Dehalobacter* increased from 2 to 17% which corresponds to a 12.5-fold change in its population. Assuming one 16S rRNA gene copy per *Dehalobacter* cell, the resulting *Dehalobacter* yield is 2.8 ± 0.05 x 10^10^ cells per mole of Cl^-^ released. This yield is two orders of magnitude lower than that reported for the population grown using 1,2,4-TCB as electron acceptor (4.6×10^12^ to 1.8×10^13^ cells per mole Cl^-^ released).^26^ The lower yield may be related to the fact that only two electron equivalents can be transferred per mole of MCB in contrast to four or six electron equivalents per mole of TCB depending on the resulting dechlorination products.

As mentioned earlier, in the bottles bioaugmented with culture II, the qPCR-determined *Dehalobacter* gene copies (Figure 3C) did not decline because the lindane-dechlorinating *Dehalobacter* population was replaced by the MCB-dechlorinating *Dehalobacter* population native to culture II. We were able to demonstrate this after a close examination of the 16S rRNA gene amplicon sequencing data. The *Dehalobacter* in culture I are represented by a single amplicon sequence variant (ASV), while *Dehalobacter* in culture II are represented by two dominant ASVs. The absolute abundance of *Dehalobacter* ASVs in cultures I and II, as well as in the active experimental bottles before and after inoculation with culture II are shown in Figure 5. The sum of the absolute abundance of all *Dehalobacter* ASVs, which were estimated from the product of the relative ASV abundance and the qPCR-determined Bacteria gene copies, closely matches the qPCR-determined abundance of *Dehalobacter* (Figure S6); this result adds confidence to our approach to estimate the absolute abundance of each ASV of interest.

**Figure 5.**
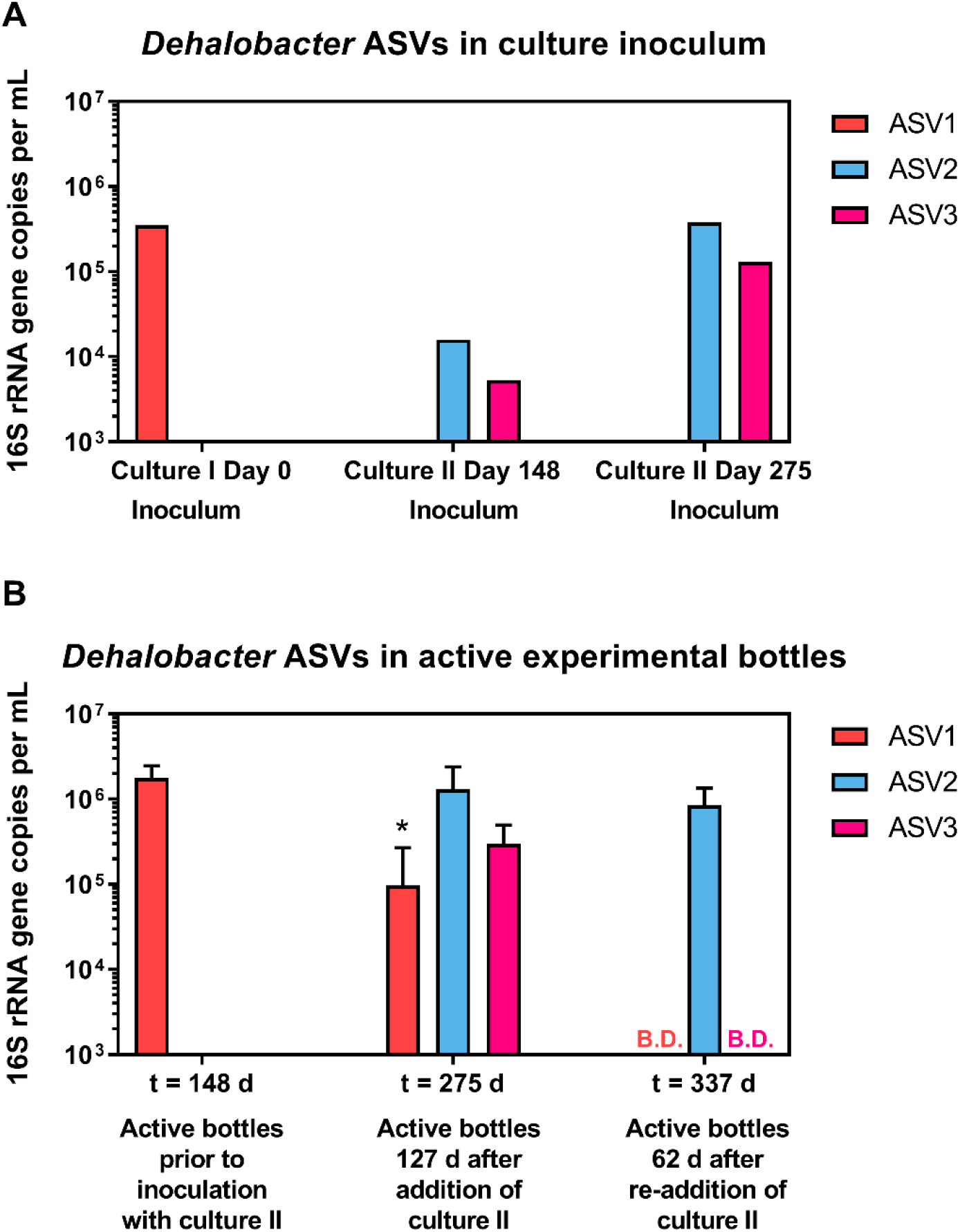
Absolute abundance of distinct *Dehalobacter* 16S rRNA amplicon sequence variants (ASVs) from culture I and culture II (A) observed in active bottles at the end of phase I and during phase II (B). Error bars represent the standard deviation of replicate experimental bottles. Absolute abundance was calculated as the product of the qPCR-determined Bacteria 16S rRNA gene copy numbers and the relative abundance of each ASV determined by 16S rRNA gene amplicon sequencing.

*Dehalobacter* ASV1 belongs to culture I, while *Dehalobacter* ASV2 and ASV3 belong to culture II (Figure 5A). There are thirteen nucleotide differences between ASV1 and ASV2, twelve differences between ASV1 and ASV3, and only one nucleotide difference between ASV2 and ASV3 (see Figure S7; ASVs can be found in the Supplementary Excel file). After addition of culture II, the population represented by ASV1 declined and was replaced by that of ASV2 and ASV3 which further confirms that the dechlorination of MCB to benzene was carried out by *Dehalobacter* populations originating from culture II. ASV1, the single dominant ASV in culture I is a 100% match to one of the *Dehalobacter* operational taxonomic units (OTU 1134628) reported in Qiao et al.^21^ for the HCH-dechlorinating culture set that includes culture I.

After the second bioaugmentation with culture II, ASV2 remained in the biaugmented bottles and ASV3 was found below detection. The differences observed in the absolute abundance of ASV2 and ASV3 in the different samples indicates that these ASVs represent different *Dehalobacter* populations in culture II. After an examination of historical 16S rRNA gene amplicon sequencing data, we were able to determine that ASV2 is in fact the dominant *Dehalobacter* ASV in the KB-1-derived enrichment cultures that dechlorinate chlorinated benzenes, i.e. this ASV was the most abundant *Dehalobacter* sequence, and representative operational taxonomic unit, in previous 16S rRNA gene amplicon analyses of the KB-1-derived chlorinated benzene-fed cultures. The 468 bp sequence of ASV2 is a 100% match to the corresponding region of the partial 16S rRNA gene (1508 bp, GI: 1221108851) of *Dehalobacter* sp. TCB1 (referred to as KB1_124TCB1 in Puentes Jácome and Edwards ^26^). Two additional ASVs, at a relative abundance lower than 0.1%, were also detected in the bottles inoculated with culture II, yet these ASVs were not consistently found among all inoculated bottles. These ASVs might represent other *Dehalobacter* populations that exist in culture II at very low abundances. The mother culture of culture II contains multiple *Dehalobacter* strains,^26, 45^ so it is not unusual to detect other ASVs that likely belong to low-abundance populations.

### Implications for HCH, MCB, and benzene contamination at field sites

In this study, we have demonstrated for the first time that lindane (γ-hexachlorocyclohexane) can be anaerobically biotransformed by sequential bioaugmentation using specialized microbial enrichment cultures. The sequential treatment resulted in the biotransformation of 73 ± 3% of the added lindane all the way to the non-toxic end products. Not only lindane, but also toxic MCB and carcinogenic benzene were biodegraded by the anaerobic enrichment cultures. Although this study focused on the γ isomer of HCH, this approach is applicable to other HCH isomers since enrichment cultures that dechlorinate α-, β-, and δ-HCH to MCB and benzene are also currently studied in the lab.^21^ The success of the different biotransformation steps in this lab scale setting was aided by the sequential nature of the treatment. Keeping the three transformation steps separate could possibly have minimized any detrimental microbial interactions between the different cultures, such as competition for the different electron donors or acceptors present. The enrichment cultures described here have been studied extensively in the lab and in-house techniques for culture growth and maintenance have improved during several years of cultivation. Specifically, benzene-degrading culture III, has recently been evaluated as an inoculum with different field site materials (soil and groundwater) and is undergoing field scale pilot tests. At the field scale, HCH remediation could be tested sequentially, analogous to the lab scale, taking into account the spatial and temporal gradients at the contaminated site and different rates of transformation. This study further establishes the key role of *Dehalobacter* spp. as chlorobenzene- and HCH-organohalide-respiring bacteria and demonstrates that sequential treatment with specialized anaerobic cultures may be explored at field sites in order to address legacy soil and groundwater contamination with HCH.

## Acknowledgements

The authors would like to thank: Shen Guo for conducting GC sampling and DNA extraction during phase III of the biotransformation experiment; Dr. Courtney Toth for performing the preliminary steps in the QIIME 2 pipeline for the analysis of the 16S rRNA amplicon sequencing data; and Suly Rambinaising for performing the qPCR runs with the Desulfuromonadales/*Geobacter* primer. L. P. J. was supported by the NSERC CREATE RENEW program [180804567], the Government of Ontario through the Ontario Graduate Scholarship program, the Ontario Research Fund INTEGRATE project [ORF-RE05-WR-01], and the Genomic Applications Partnership Program GAPP [OGI-102].

## Notes

### Competing Interest Statement

The authors have declared no competing interest.

### Summary of Updates

Oct. 26th manuscript submission to bioRxiv contained the "Error! Reference source not found" messages throughout. This has been fixed

