## Supplemental information for "Biotransformation of lindane (γ-hexachlorocyclohexane) to non-toxic end products by sequential treatment with three mixed anaerobic microbial cultures"

Figure S5. Coupling of MCB-to-benzene-dechlorination to growth of a *Dehalobacter* population in the scaled-up culture II (500 mL). Cumulative benzene production from MCB is shown in A. 16S rRNA gene copies per mL of culture of *Dehalobacter* (*Dhb*) and Bacteria (Bac) measured by qPCR are shown in B. In B, the error bars represent the range of qPCR technical duplicates. The % above the green *Dhb* bars indicates the calculated relative abundance of *Dehalobacter* at each time point, i.e. 100*(16S rRNA gene copies of *Dehalobacter*/16S rRNA gene copies of Bacteria). ........ 6

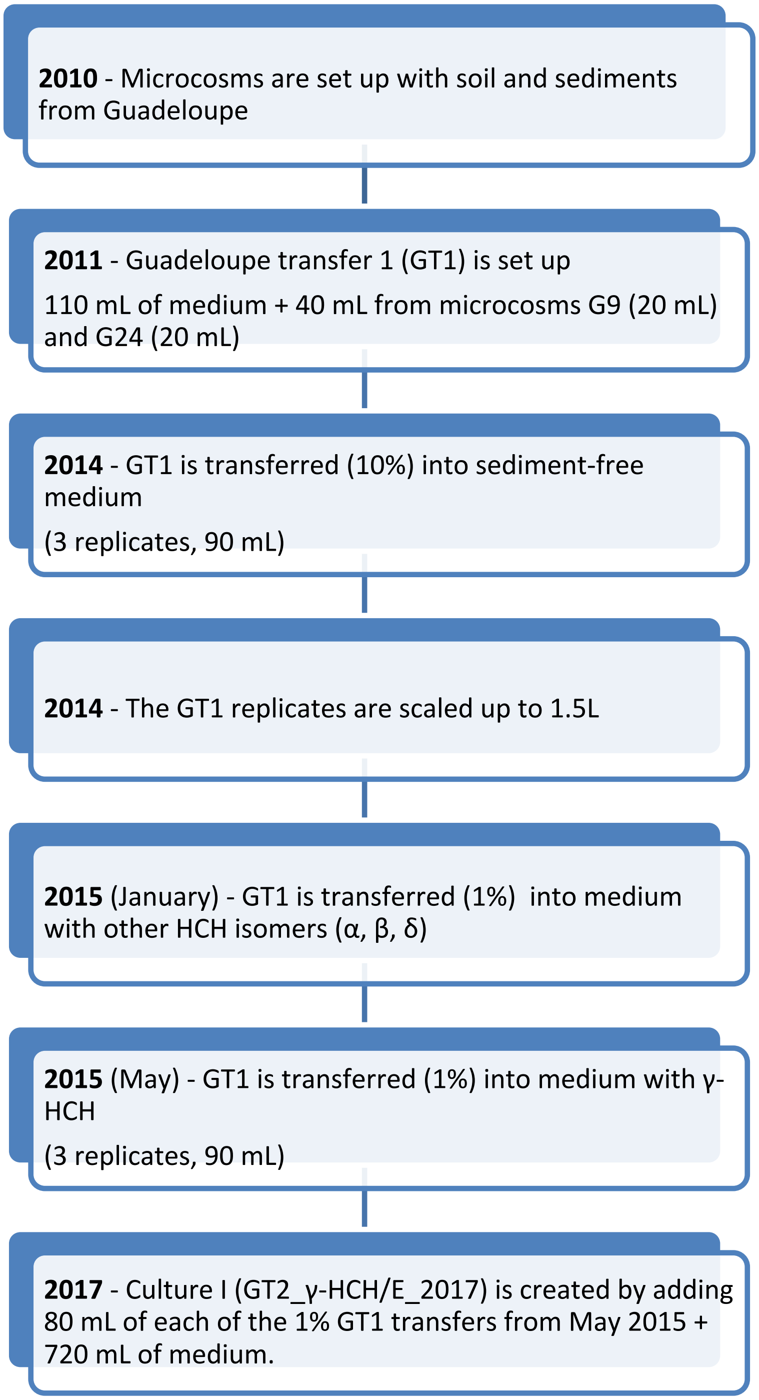

Figure S1. History of Culture I, a microbial enrichment culture that dechlorinates lindane (γ-hexachlorocyclohexane) into monochlorobenzene and benzene. From 2010 to 2015, the cultures were fed a combination of acetone and ethanol as electron donor. In 2016 and 2017, the ratio of added electron donor (ethanol only) was reduced to 10 to 1, and 5 to 1, electron equivalents of donor to acceptor, respectively.

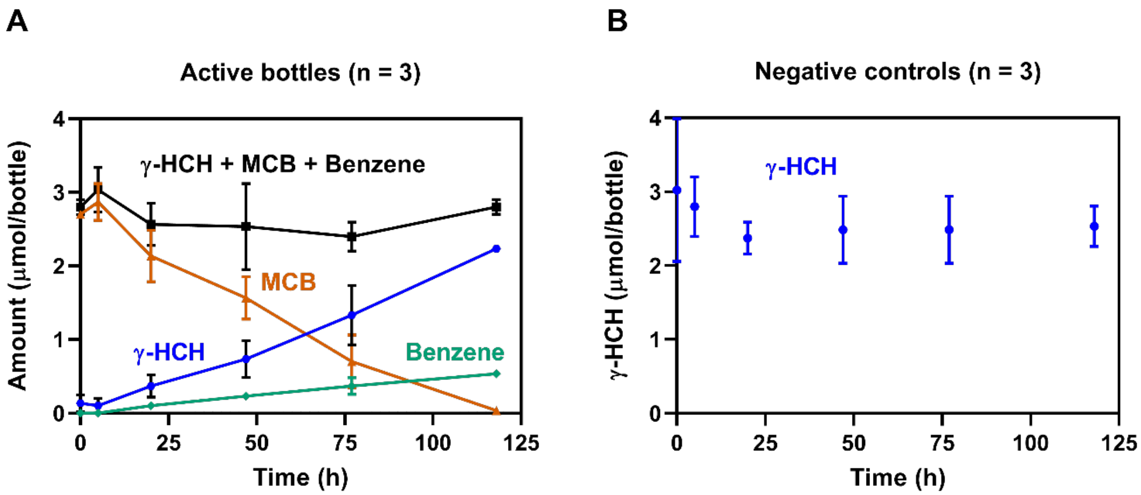

Figure S2. Dechlorination of γ-hexachlorocyclohexane (γ-HCH) to MCB and benzene by culture I. Replicate experimental bottles (n = 3) are shown in A and negative controls (n = 3) are shown in B. This figure was first published as part of the supplementary information in Qiao, et al. [1]

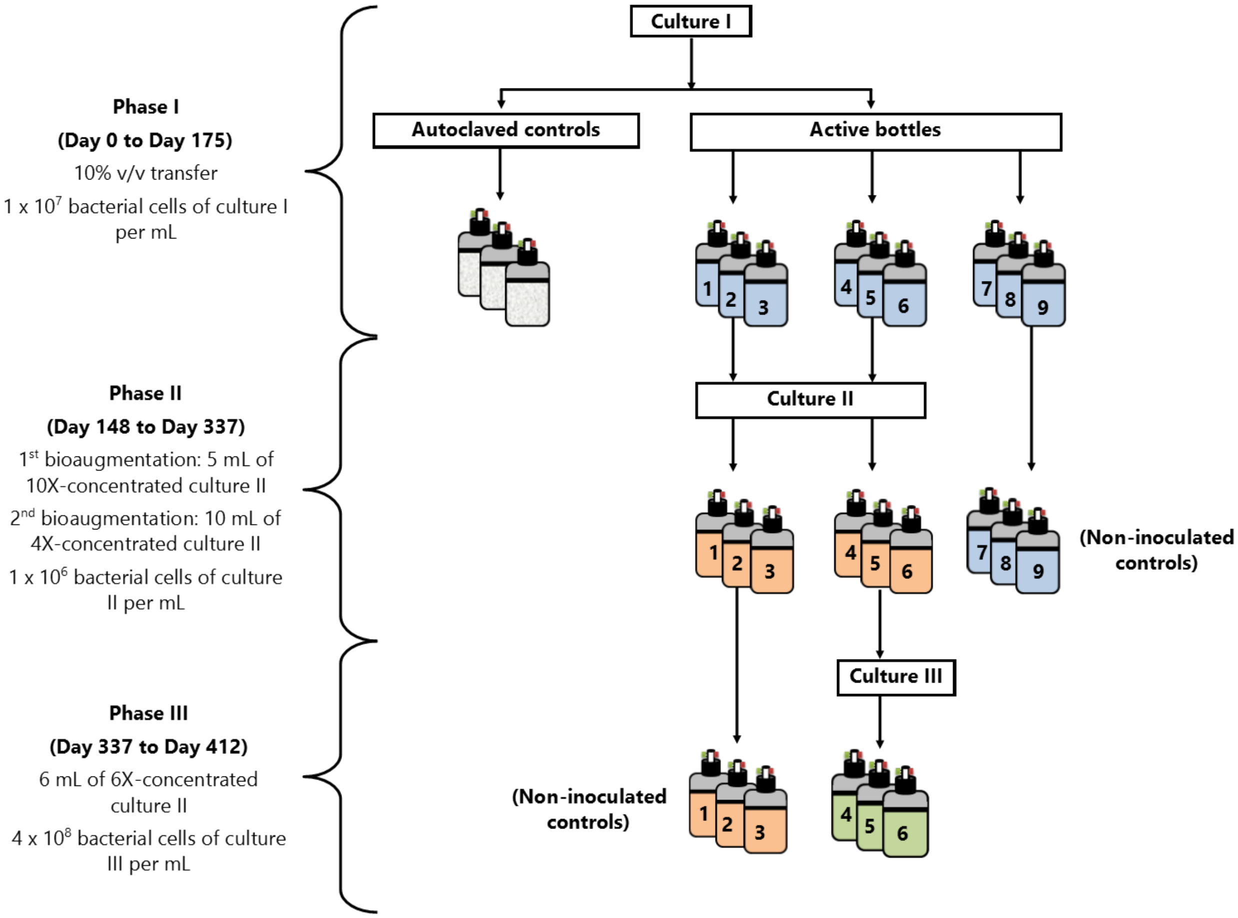

Figure S3. Schematic of the different phases of the sequential lindane (γ-hexachlorocyclohexane) biotransformation experiment. An amount equivalent to 3.8 mg of γ-HCH was added to each bottle during phase I (150 µmol/L of culture, 13.5 µmol/bottle). Each experimental bottle had a working volume of 90 mL. Experimental bottles, labelled 1 to 9, are color-coded according to each sequential biotransformation phase. Inoculum amounts added to active bottles during each phase are shown on the left.

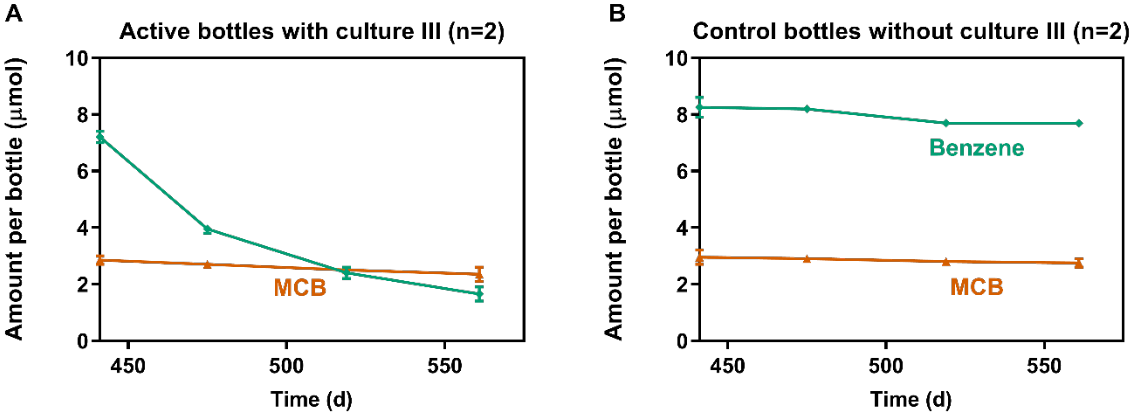

Figure S4. Benzene degradation in active bottles (A) after exogenous benzene addition (neat benzene was added to the bottles directly) to confirm that culture III remained active. MCB and benzene concentration in control bottles is shown in B. Error bars represent the range of measurements in duplicate experimental bottles (n = 2).

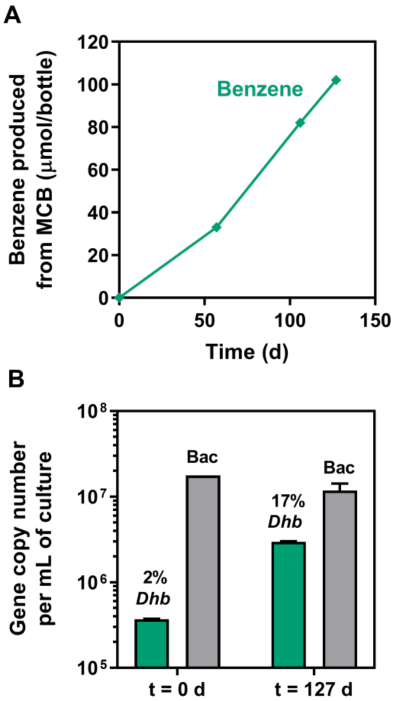

Figure S5. Coupling of MCB-to-benzene-dechlorination to growth of a *Dehalobacter* population in the scaled-up culture II (500 mL). Cumulative benzene production from MCB is shown in A. 16S rRNA gene copies per mL of culture of *Dehalobacter* (*Dhb*) and Bacteria (Bac) measured by qPCR are shown in B. In B, the error bars represent the range of qPCR technical duplicates. The % above the green *Dhb* bars indicates the calculated relative abundance of *Dehalobacter* at each time point, i.e. 100*(16S rRNA gene copies of *Dehalobacter*/16S rRNA gene copies of Bacteria).

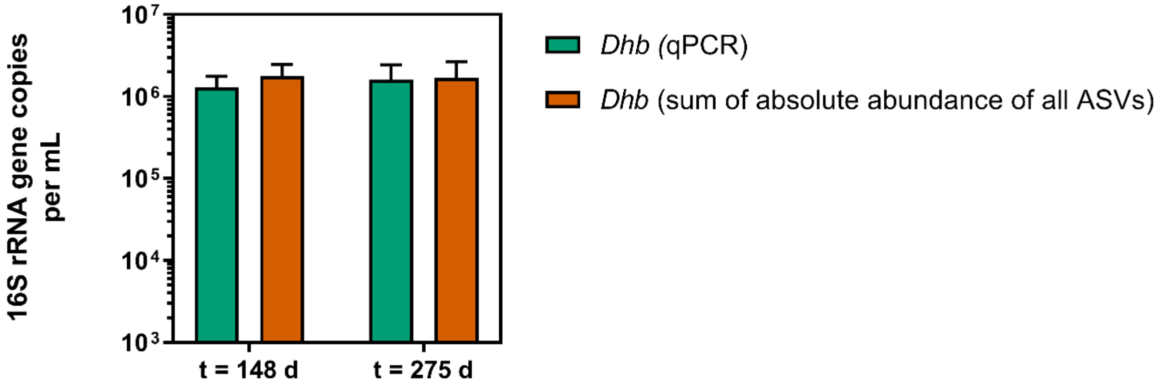

Figure S6. Absolute abundance of *Dehalobacter* (*Dhb*) in culture II-bioaugmented bottles determined by qPCR (green bars) and by the sum of the absolute abundance of all *Dhb* ASVs estimated as the product of the qPCR-determined Bacteria 16S rRNA gene copy numbers and the relative abundance of each ASV determined by 16S rRNA gene amplicon sequencing (orange bars).

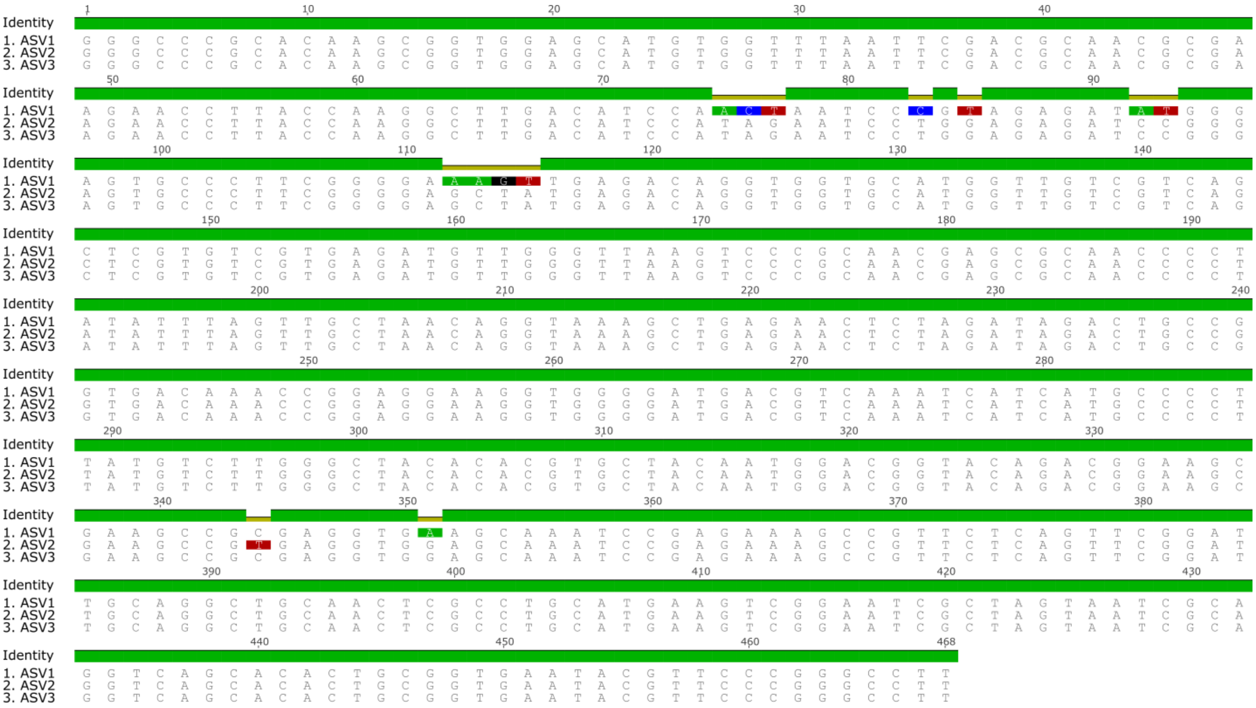

Figure S7. Multiple sequence alignment of *Dehalobacter* amplicon sequence variants (ASVs) identified in culture I (ASV1) and culture II (ASV2 and ASV3).

Table S1. Primer sequences used for qPCR during the lindane biotransformation study.

| Target | Primer short name | Primer Sequence 5' - 3' | Expected amplicon length (bp) | Primer set specificity[a] | Reference |
|---|---|---|---|---|---|
| *Dehalobacter* | Dhb 477f | GATTGACGGTACCTAACGAGG | 201 | 82/3,196,041 Kingdom Bacteria<br>82/167 Genus *Dehalobacter* | Grostern and Edwards [2] |
| | Dhb 647r | TACAGTTTCCAATGCTTTACG | | | |
| *Desulforomonadales/ Geobacter* | Geo_67r | GCAGAACCTTACCTGGGCTT | 213 | 2,105/3,196,041 Kingdom Bacteria<br>1,193/9,614 Order Desulforomonadales<br>1,179/6,574 Genus *Geobacter* | This study. |
| | Geo_279r | CACCTTCCTCCGGTTTGACA | | | |
| General Bacteria | Bac 1055f_Dhb | ATGG**Y**TGTCGTCAGC T | 338 | 875,618/3,196,041 Kingdom Bacteria | Ferris, et al. [3]<br>Note that the forward primer used in the present study has a Y at the fourth nucleotide position. |
| | Bac 1392 | ACGGGCGGTGTGTAC | | | |

[a]16S rRNA primer specificity was determined using the Ribosomal Database Project (RDP) Probe Match tool (https://rdp.cme.msu.edu/probematch/search.jsp, accessed September 28, 2020); no mismatches were allowed. The values shown indicate the number of sequences captured by the primer set relative to the total number of available sequences for a given taxonomy.

Table S2. Calibration information for the qPCR assays performed during the lindane biotransformation study.

| qPCR Target | Phase | Slope | y-intercept | $R^2$ | Efficiency | Limit of detection (copies $\mu l^{-1}$) |
|---|---|---|---|---|---|---|
| General Bacteria | I/II | -3.537 | 35.444 | 0.984 | 92% | 5.3E+02 |
| *Dehalobacter* | I/II | -3.589 | 36.250 | 0.998 | 90% | 5.3E+02 |
| *Desulforomonadales/Geobacter* | I/II | -3.699 | 37.715 | 0.999 | 86% | 1.9E+02 |
| General Bacteria | III | -3.364 | 35.299 | 0.996 | 98% | 2.2E+02 |

**Supplementary Materials and Methods**

**i. Concentration of culture inoculum for phase I and phase II of the biotransformation experiment**

Culture II was concentrated by centrifugation at 16,000x g for 30 min at room temperature using screw-capped Nalgene® polypropylene bottles fitted with O-rings and sealed with anaerobic tape. The culture supernatant was removed and used to re-suspend the pellet to the desired final volume. Culture III was concentrated using a two-step centrifugation cycle at room temperature. First, the culture was centrifuged at 6,000x g for 10 min using the swing-bucket rotor. Then, the culture was centrifuged at 8,000x g using a fixed-angle rotor. The culture supernatant was removed and subsequently used to re-suspend the pellet to the desired final volume.

**ii. Analysis of monochlorobenzene (MCB), benzene, and methane by GC-FID**

Aqueous samples (1 mL) were added to 5 mL of acidified (using 6N HCl) deionized water (pH < 2). Samples were then equilibrated in an Agilent G1888 autosampler at 70°C for 40 min. After equilibration, 3 mL of headspace sample were injected into an Agilent 7890A GC equipped with an Agilent GS-Q plot column (30 m length, 0.53 mm diameter) via a packed inlet. The carrier gas was helium at a flow rate of 11 mL $min^{-1}$. The temperature of the injector and the detector were set at 200°C and 250°C, respectively. The oven was held at 35 °C for 1.5 min, ramped to 100°C at a rate of 15°C per min, ramped to 185°C at a rate of 5°C per min, held for 10 min, ramped to 200°C at a rate of 20°C per min, and held at 200°C for 10 min.

**iii. Analysis of γ-hexachlorocyclohexane (lindane) performed for the data shown in Figure S2.**

Lindane was quantified based on a previously published protocol[4] with some modifications described as follows. Briefly, 250 µl of an aqueous sample was added to a 2 mL screw-cap autosampler vial. Hexane (500 µL) was added to the same vial as the extraction solvent. The samples were placed in a vortex shaker for 2 min. The hexane extract (supernatant) was then transferred via a Pasteur pipette to a new 2 mL autosampler vial and the procedure was repeated once in order to obtain a combined ~ 1 mL extract. This extract (1 µL) was analyzed by gas chromatography coupled with mass spectrometry (GC-MS) using a Varian Ion trap equipment with a 5% phenyl 95% methylpolysiloxane (VF-5MS) column (length: 30 m, width: 0.25mm, film thickness 0.25 µm). The temperature of the injector was set to 250°C and a split ratio of 25:1 was used. The oven was held at 75°C for 0.5 min, ramped to 100°C at a rate of 25°C per min, held for 1 min, ramped to 180°C at a rate of 25°C per min, held for 1 min, ramped to 300°C at a rate of 25°C per min, and finally held at 300°C for 2 min. Helium was used as the carrier gas at a flow rate of 1 mL per min. An external calibration curve was obtained using standards prepared in isooctane (lindane concentrations ranged from 0.2 to 10 mg/L).

### iv. qPCR protocol and quantification of 16S rRNA gene copies per mL of culture

The qPCR prep was conducted in a UV-treated PCR cabinet (ESCO Technologies, Hatboro, PA) with the fan on. Each 20 ul qPCR reaction contained 10 µl of 2×SsoFast™ EvaGreen® (Bio-Rad, Hercules, CA), forward and reverse primers (0.5 µM each), and 2 µl of template DNA. The amplification program included an initial denaturation step at 98°C for 2 min, followed by 39 cycles of 5 s at 98°C and 10 s at the corresponding annealing temperature for each primer set. Quantification of total bacteria and *Dehalobacter* was performed using 10-fold serial dilutions of plasmid DNA (containing a nearly full-length 16S rRNA *Dehalobacter* sequence) as standards. Quantification of Desulforomonadales/*Geobacter* was performed using 10-fold serial dilutions of purified PCR product amplified from HCH-dechlorinating enrichment cultures. The qPCR reactions were conducted using a BIO-RAD CFX96 Touch™ Real-Time PCR Detection System and the CFX Manager software. The number of 16S rRNA gene copies per mL of culture was calculated assuming a 100% DNA extraction efficiency and taking into account the DNA dilution and culture volumes used for DNA extraction.
